# Protocol for the production of an Arenavirus and Hantavirus host-pathogen database: Project ArHa

**DOI:** 10.1101/2025.01.17.633514

**Authors:** David Simons, Ricardo Rivero, Ana Martinez-Checa Guiote, Harry Gordon, Gregory C. Milne, Grant Rickard, David W. Redding, Stephanie N. Seifert

## Abstract

1

Arenaviruses and Hantaviruses, primarily hosted by rodents and shrews, represent significant public health threats due to their potential for zoonotic spillover into human populations. Despite their global distribution, the full impact of these viruses on human health remains poorly understood, particularly in regions like Africa, where data is sparse. Both virus families continue to emerge, with pathogen evolution and spillover driven by anthropogenic factors such as land use change, climate change, and biodiversity loss. Recent research highlights the complex interactions between ecological dynamics, host species, and environmental factors in shaping the risk of pathogen transmission and spillover. This underscores the need for integrated ecological and genomic approaches to better understand these zoonotic diseases. A comprehensive, spatially and temporally explicit dataset, incorporating host-pathogen dynamics and human disease data, is crucial for improving risk assessments, enhancing disease surveillance, and guiding public health interventions. Such a dataset (ArHa) would also support predictive modelling efforts aimed at mitigating future spillover events. This paper proposes the development of this unified database for small-mammal hosts of Arenaviruses and Hantaviruses, identifying gaps in current research and promoting a more comprehensive understanding of pathogen prevalence, spillover risk, and viral evolution.

**Strengths and Limitations of this study:**

- This dataset combines detailed spatial and temporal information, providing a unique resource for understanding geographic and temporal trends in Arenavirus and Hantavirus host-pathogen relationships.
- By explicitly quantifying sampling biases and detection efforts, the dataset allows more robust and accurate asssessments of pathogen prevalence and distribution.
- The dataset offers a platform for linking ecological data with human health outcomes, supporting the identification of spillover hotspots.
- The dataset relies on published material, which may vary in terms of detail, accuracy and completeness. Missing or imprecise information may limit the reliability of subsequent analyses.
- The dataset will be produced as a static resource which could limit its relevance over time as emerging data will not be added.

## 3 Introduction

Arenaviruses and Hantaviruses are globally distributed pathogens which primarily infect rodents (order Rodentia) and shrews (order Eulipotyphla) [1,2]. Some of these Arenaviruses and Hantaviruses occasionally spillover (cross-species transmission of a parasite into a host population not previously infected) into human populations, with variable morbidity and mortality rates [3]. However, the overall human health impact remains poorly understood in many cases [4,5]. Arenaviruses, include *Mammarenavirus lassaense* (LASV), responsible for Lassa fever in West Africa, and *Mammarenavirus juninense* (JUNV), which causes Argentine haemorrhagic fever in Argentina [6,7]. Lassa fever is estimated to infect 900,000 individuals annually across West Africa, with over 200 deaths reported from Nigeria in 2024 [8,9]. In contrast to the limited distribution of LASV, *Mammarenavirus choriomeningitidis* (causing Lymphocytic choriomeningitis) has a global distribution with few infections reported [10].

Hantaviruses, including *Orthohantavirus puumalaense* (PUUV) and *Orthohantavirus seoulense* (SEOV) cause haemorrhagic fever with renal syndrome (HFRS) in Europe and Asia, while *Orthohantavirus sinnombreense* (SNV) causes hantavirus pulmonary syndrome (HPS) in the Americas [11–13]. HFRS is associated with 23,000 reported annual infections, but fewer than 100 deaths, whereas HPS, which has a higher case fatality rate (12-45%) is reported in fewer than 300 cases annually [14]. The distribution and human health impact of hantaviruses in Africa is even more poorly understood [15].

Recent research indicates that both Arenaviruses and Hantaviruses continue to emerge, with pathogen evolution and spillover driven by anthropogenic factors such as land use change, climate change and small-mammal biodiversity loss [16,17]. In addition to direct biodiversity changes, the structure and dynamics of host populations and their community interactions have also been shown to influence pathogen prevalence [18,19]. Human activities such as deforestation, urbanization, and agricultural expansion bring humans into closer (e.g., increased frequency and intensity) contact with rodent reservoirs, increasing the likelihood of spillover events [20,21]. Changes in climate, land use, rodent ecology, and human behaviour all affect human-reservoir contact, contributing to the complex human-animal-environment nexus that drive pathogen spillover and persistence [22–24]. Although human-to-human transmission is a concern (e.g., Disease X), most infections result from rodent-to-human transmission, with LASV classified as a priority pathogen by the WHO due to its potential for human-to-human spread [4,25].

Understanding the economic and societal impacts of these diseases is also critical. In addition to direct healthcare costs, there are significant indirect costs (e.g., long-term productivity losses), which exacerbate economic strain on vulnerable populations [4]. Outbreaks of emerging rodent-borne zoonoses, like HPS during the Four Corners Outbreak (1993) or Yosemite outbreak (2012), demonstrate the broader social and economic disruptions that these diseases can cause [26,27]. Long-term health consequences, such as sensorineural hearing defects in survivors of Lassa fever, further complicate the disease burden [28]. Although only ∼20% of LASV infections are symptomatic, the full scale of both acute and long-term disease remains poorly understood [29]. Under-reporting of these diseases, likely due to difficulties in diagnosing acute infections, means that many cases go undetected, particularly in endemic regions [14].

The traditional paradigm in which a specific pathogen is hosted by a single reservoir species is being overturned by increased pathogen surveillance in endemic settings [30]. These data have identified multiple potential hosts for pathogens including LASV, SEOV and SNV [13,31,32]. Similarly, a single host species may be a host of multiple Arenaviruses and Hantaviruses throughout its geographic distribution, for example *Mastomys natalensis*, the primary host of LASV is known to host at least 7 distinct Arenaviruses [33].

The ecology of rodent hosts — such as population dynamics, behaviour, habitat preferences, and community interactions — directly influences pathogen transmission and spillover risk [34]. Linking ecological and genomic data enables the identification of genetic markers associated with increased virulence or transmissibility, providing critical insights into the evolutionary trajectories of Arenaviruses and Hantaviruses [35,36]. Small-mammal sampling is essential for uncovering how these ecological factors moderate the prevalence, persistence, and spread of zoonotic pathogens within their communities [37,38]. For example, fluctuations in host population density or shifts in community composition, often driven by habitat changes or seasonal cycles, can amplify transmission within reservoirs or increase human exposure risk [39,40]. Understanding interactions between host species within shared environments, including competition or co-occurrence, further elucidates how pathogens circulate, expand, and evolve across interconnected populations [41–43]. These insights underscore the value of systematic small-mammal sampling to inform surveillance, predict areas of heightened spillover risk, and develop targeted strategies to mitigate zoonotic disease emergence.

A dataset that includes detailed spatial and temporal data on rodent hosts and their pathogens would enable the linkage of local ecological findings with public health patterns at broader scales. While human disease data are often aggregated at national or subnational levels to preserve patient anonymity, clinical practitioners and public health authorities will have access to higher-resolution case data. For these stakeholders, integrating ecological data with localised human disease patterns could enhance the identification of hotspots of transmission risk and inform more targeted interventions. Data fragmentation currently limits efforts to connect local-scale, high-intensity research on host-pathogen dynamics to broader-scale trends, which impairs predictive modelling and disease management efforts. A unified dataset would support these efforts, enabling better understanding of ecological drivers of zoonotic disease risk, viral evolution, and pathogen emergence across landscapes and over time, improving both risk assessments and public health interventions [44].

Historically, research on rodent-borne zoonoses has been uneven, with undersampling in high-risk regions like Africa, South Asia, and Southeast Asia [31,45,46]. Identifying these undersampled regions is crucial for guiding future research and prioritizing viral discovery efforts [47]. Addressing these gaps will improve our understanding of pathogen diversity, distribution, and evolution, and help predict and mitigate future spillover events [48,49]. A comprehensive global sampling effort will highlight knowledge gaps and inform future research priorities.

A unified and comprehensive dataset on Arenaviruses and Hantaviruses in wild-caught small mammals is urgently needed to address these gaps. By collating spatial, temporal, and genomic data, such a dataset will be a critical resource for identifying undersampled regions and potential hosts, guiding viral discovery, and informing risk assessments, predictive models, and public health interventions. Adhering to Open Science principles, we will ensure that research tools, data extraction methods, and processing code are shared on suitable platforms following the FAIR guidelines [50]. This will ensure global accessibility and foster evidence-based decision-making and collaboration in disease surveillance. As automated tools increasingly drive scientific research, we must standardize datasets to avoid overlooking valuable data. By curating this dataset, we aim to preserve scientific knowledge and ensure its accessibility to global researchers through platforms like the Global Biodiversity Information Facility (GBIF) [51].

Existing data on rodent-pathogen associations are often limited by geographic and temporal constraints, which limits broader analyses of pathogen prevalence, spillover risk, and range expansion. The proposed ArHa database will address these limitations by synthesizing and standardizing data from diverse sources, enabling research on host-pathogen dynamics, range expansion, and spillover risk. This unified resource will strengthen predictive models of zoonotic diseases, improve public health strategies, and enhance preparedness for future disease threats.

## 4 Method

### 4.1 Search strategy

We searched NCBI PubMed and Clarivate Web of Science for relevant citations (see Table 1 for search terms), with no restrictions on publication date, language, or sampling locations. The search terms used are shown in Table 1, which details the search terms and results from both databases.

**Table 1:** Search terms used for identifying relevant literature.

| Search term | PubMed | Web of Science |
| --- | --- | --- |
| 1. rodent* OR rat OR mouse | 3,878,308 | 4,090,059 |
| 2. shrew OR eulipotyphla | 7,582 | 9,108 |
| 3. arenavir* | 2,525 | 1,966 |
| 4. hantavir* | 4,316 | 5,084 |
| 5. 1 OR 2 | 3,883,403 | 4,096,018 |
| 6. 3 OR 4 | 6,746 | 6,899 |
| 7. 5 AND 6 | 3,100 | 2,904 |

Search terms 1-4 (Table 1) included expanded terms. Citations returned by the searches were downloaded and de- duplicated by matching on titles, authors, publication identifiers, or digital object identifiers. The initial search (conducted 2023-10-16) resulted in 2,755 unique publications. These citations were uploaded to the Rayyan platform to assess against inclusion and exclusion criteria [52].

### 4.2 Study screening

#### 4.2.1 Inclusion criteria

We screened studies against the following inclusion criteria. Studies were included if they reported:

1. Rodent OR Shrew sampling from wild populations AND
2. Direct or indirect detection of microorganisms in the Arenaviridae and Hantaviridae families AND
3. Information on the sampling location of small mammals

#### 4.2.2 Exclusion criteria

Studies were excluded if they:

1. Did not contain information on the host species of the sample for direct or indirect detection of Arenaviridae or Hantaviridae,
2. Reported experimental infections or solely human infections,
3. Were abstracts or conference proceedings that did not provide any description of the method of animal sampling or microorganism detection.

Direct detection was defined as detection via nucleic acid amplification tests (e.g., Polymerase Chain Reaction (PCR)) or virus culture, indirect detection was defined as detection of specific antibodies or antigens.

We first screened the study titles and abstracts. For articles published in languages other than English we used automated translation services (e.g., Google Translate) if study authors were unable to review the manuscript in its published language. All titles and abstracts were then assessed against our inclusion and exclusion criteria. Assessments were made by two members of the study team (DS and RR). Studies deemed to meet all inclusion criteria by at least one author proceeded to full-text review.

Reference chaining was performed on studies considered for inclusion at the full text screening stage and relevant publications were added as manually identified relevant articles. The PRISMA flow chart for the search process is shown in Figure 1 which also indicates the status of *pilot* data extraction (10% of the 917 studies included at title and abstract stage). The search will be re-run to capture more recently published articles when data extraction from the current search has been completed.

**Figure 1:**
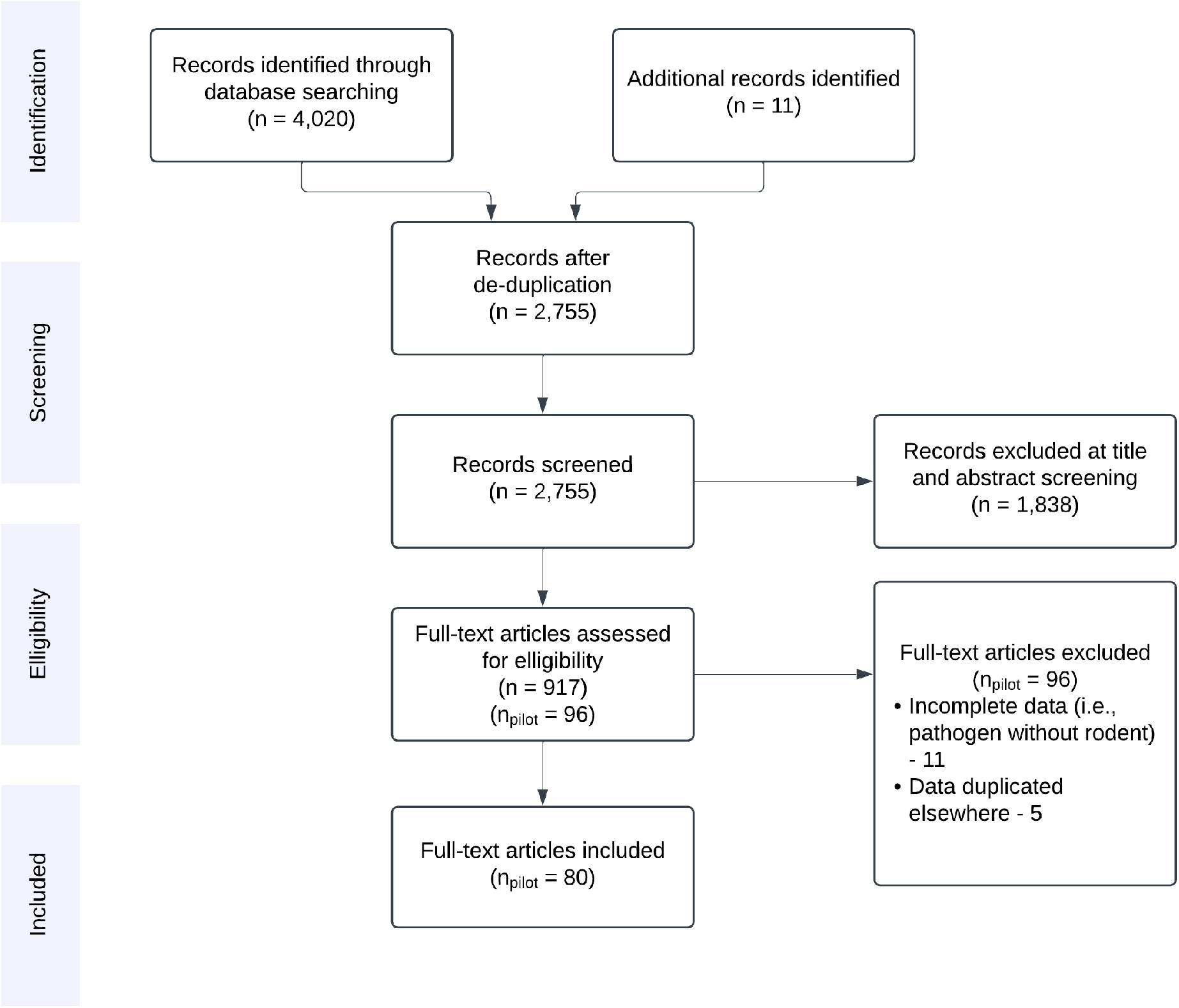
PRISMA flow diagram for records identified in the initial search conducted 2023-10-16. Studies excluded at full text review included those with incomplete data on 1 of pathogen, host, or study location. For studies containing duplicated data, the highest resolution dataset (i.e., temporal, spatial, taxonomic) was retained.

### 4.3 Data extraction

We aimed to extract included study meta-data and to produce three linked datasets that could be used to assess sampling of a) small-mammal hosts, b) viral prevalence and c) genetic variability. We have developed and refined the current data extraction tools in the pilot search (Figure 1). A publicly available RShiny web-application will be developed to present the extracted data while the search is ongoing [53].

#### 4.3.1 Included studies

Included study meta-data was abstracted to ensure appropriate attribution for each rodent, pathogen and genomic material record to the original publication that presented it (Table 2). Each included study was assigned a unique identifier which provides linkage to the reference. We extracted reported sampling effort, when this information was available (e.g., number of trap-nights). For articles not reporting sampling effort at the study level we inferred total sampling effort by summing effort across individual trapping sessions and sites. For studies that did not report a measure of sampling effort amenable to imputation (e.g., a variable number of traps per trapping line, or days per study session) we recorded a value of ‘not reported’. Finally, we recorded the level of data aggregation of rodent or pathogen sampling, classified as individual (data at the individual level) or summarised (data were aggregated; e.g., at site or sampling session level).

**Table 2:** Study information extraction sheet

| Column name | Description |
| --- | --- |
| study_id | An internal unique identifier for the included study |
| full_text_id | The internal ID for the full text manuscript and its associated citation |
| datasetName | The title of the manuscript, report or book section |
| sampling_effort | A free text entry to capture the effort of sampling, ideally in trap nights for rodent studies |
| data_access | Whether data is available for individual small mammals or whether it is aggregated |
| data_resolution | The level of aggregation available |
| linked_manuscripts | The DOI or weblink to other studies including the same dataset either in its entirety or a subset, this will be used to attempt to de-duplicate data |
| notes | Details that may be important for interpreting the extracted data |

#### 4.3.2 Small-mammal sampling

Data were extracted into a rodent sheet (Table 3) at the highest available level of temporal and spatial resolution. For example, a study reporting a single species at four spatially distinct sampling sites would be associated with four records. If this same study reported observations across four distinct trapping sessions at each of these sites there would be 16 (4^2^) records associated with the study. Studies presenting data on *N* individuals are associated with *N* records. These records may not be for individuals (as would be the case for capture-mark-recapture studies), for example, they may be the same individual detected over multiple sessions. For this reason, we do not report a unique identifier at the individual level.

**Table 3:** Extracted rodent sampling variables

| Column name | Description |
| --- | --- |
| rodent_record_id | A unique internal identifier for the rodent record. Record resolution is depends on the extent of aggregation in the study |
| study_id | A unique identifier to link a record to its respective study |
| scientificName | The binomial species name of the small-mammal reported in the study |
| eventDate | The period in which sampling was conducted |
| locality | The location of sampling effort, reported to the highest spatial resolution available |
| country | Country where trapping occurred. For multinational studies where counts are not disaggregated by country, all countries are reported |
| verbatimLocality | High level habitat type will be recorded here at the scale for which trapping is recorded, particularly useful when locality does not discriminate between multiple sampling sites |
| coordinate_resolution | The spatial resolution of coordinates |
| decimalLatitude | Latitude in decimal format, converted from reported coordinates as required |
| decimalLongitude | Longitude in decimal format, converted from reported coordinates as required |
| individualCount | The number of detected individuals associated with a record |
| trapEffort | Trap effort (recorded as number of trap nights) associated with a record |
| trapEffortResolution | The resolution of trapping effort for the record |

Identification of a detected small mammal was recorded as reported within the article, we will not systematically assess the method of identification and therefore assume accurate identification as reported within the articles. Where identification is to genus or higher taxonomic level we report to this level (i.e., *Rattus spp*., Rodentia). Further details are provided below on data processing and species name harmonisation.

Since we expected studies to report sampling location to varying degrees of resolution we extract `locality` as the highest resolution of location data that could be matched to administrative levels within a country (e.g., city, county). In cases where location is reported to a higher spatial resolution which may not map to administrative levels (e.g., ‘Site A’) we also include `verbatimLocality`. For studies not reporting geographic coordinates of sampling but including some information that describes the location (i.e., the name of the village sampled) we will locate coordinates through several methods. Locations will be searched for in Google Maps, Wikipedia or the Geographic Names Server provided by the National Geospatial-Intelligence Agency USA.

#### 4.3.3 Pathogen sampling

The pathogen sheet (Table 4) reports the assays used to detect Arenaviruses and Hantaviruses. Analogously to the rodent sheet, a record is produced at the highest resolution of pathogen sampling. One-to-many matching was permitted to record all data in cases where a single rodent was tested for a pathogen using multiple assays (e.g., serology and PCR).

**Table 4:**
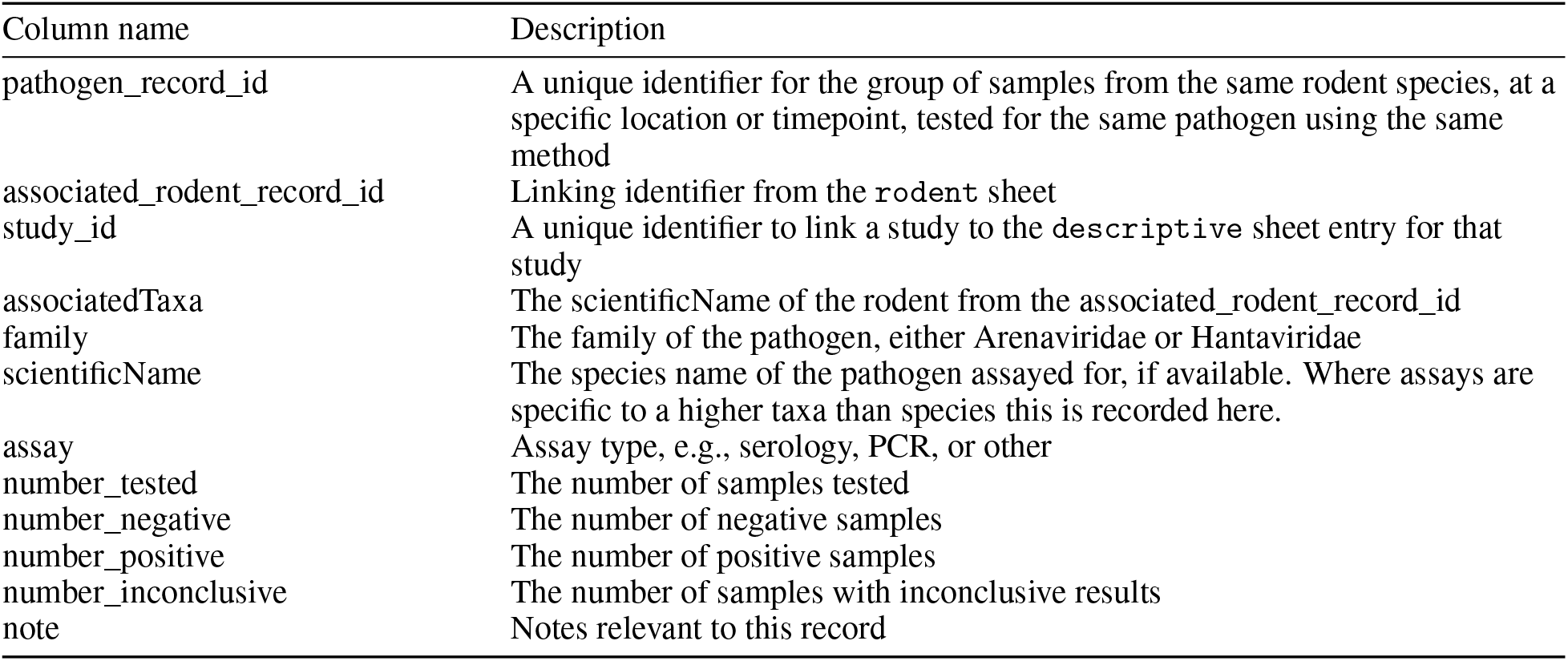
Pathogen information extraction sheet

The species or family targeted by the assay was recorded as reported within the relevant article; we did not make any assessment of the suitability, sensitivity or specificity of an assay. Where authors describe an assay as being family-wide (e.g., hantavirus antibody detection) we retain this level of specificity in the pathogen record. We recorded the number of samples tested using the assay for a specific pathogen. This may differ from the number of counted small-mammals as not all samples may not have been suitable for testing, or the study authors may have decided to subset available samples. The number of positive and negative samples are related to this tested number. Where reported we also extracted the number of samples with an inconclusive test result. Similarly to above, we recorded positives and negatives as reported by study authors and make no assessment of these classifications.

#### 4.3.4 Sequences

If studies include complete or partial nucleotide sequences of hosts or viruses archived in NCBI (USA), EMBL (Europe), DDBJ (Japan) or CNGBdb (China) they will be linked through the sequences sheet (Table 5). A record will be produced for each accession available. A sequence_record_id will be associated with each accession and depending on whether the sequence relates to a pathogen or host each record will be associated with one or both of these (i.e., a pathogen sequence will be linked to both a `pathogen_record_id` and `rodent_record_id`, while a host sequence will only be linked to a `rodent_record_id`). Many-to-one matching of sequences may occur for several reasons, first, Hantaviruses (three) and Arenaviruses (two) contain different number of genome segments and not all may have been successfully sequenced for each acutely infected small mammal. Second, reporting of pathogen sampling may be aggregated with multiple sequences obtained from a single reported assay (e.g., pooled sampling). Pathogen sequences obtained from infected humans will be extracted but will only be linked at the study level if reported within the manuscript from the same geographic location or temporal period.

**Table 5:**
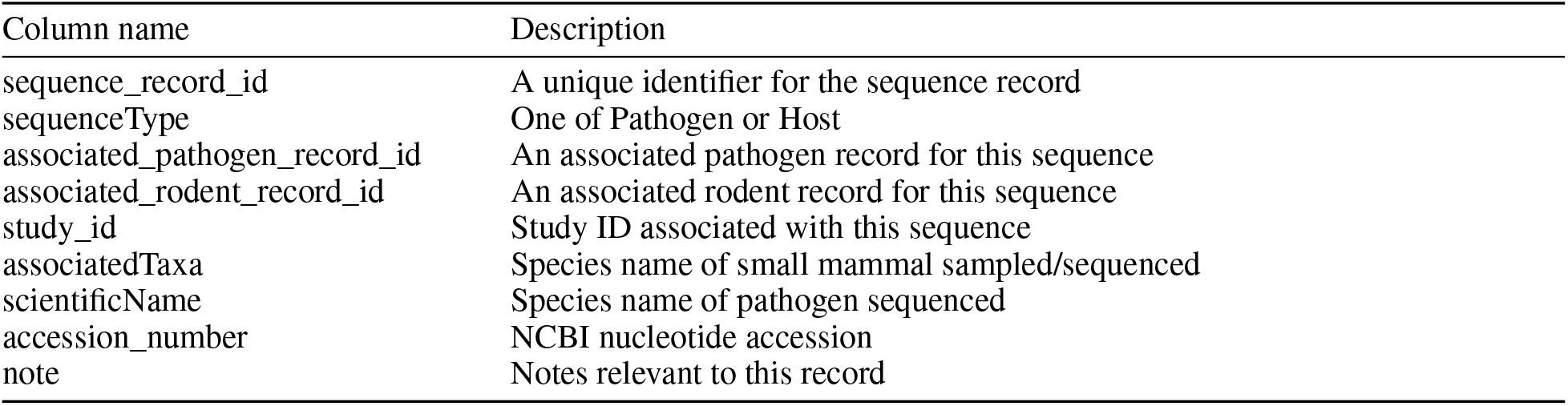
Pathogen sequences extraction sheet

| Column name | Description |
| --- | --- |
| sequence_record_id | A unique identifier for the sequence record |
| sequenceType | One of Pathogen or Host |
| associated_pathogen_record_id | An associated pathogen record for this sequence |
| associated_rodent_record_id | An associated rodent record for this sequence |
| study_id | Study ID associated with this sequence |
| associatedTaxa | Species name of small mammal sampled/sequenced |
| scientificName | Species name of pathogen sequenced |
| accession_number | NCBI nucleotide accession |
| note | Notes relevant to this record |

### 4.4 Data processing

This section describes data cleaning and harmonisation. All data processing will be conducted in R with scripts retained in a version controlled git repository [54,55]. Raw data will be downloaded from Google Sheets using the `googledrive` API in R, with date stamped files stored locally [56].

#### 4.4.1 Species name harmonisation

To standardise reported species we will match reported names to both the GBIF backbone taxonomy and the NCBI taxonomy database [51,57]. The `taxize R` package was used to query the APIs of these platforms [58]. Unmatched names are corrected to an accepted matching name if the species has been identified as a subspecies or the genus name has changed, or allocated to a higher taxa if there is no suitable matching at the species level (e.g., “*Mus/Nannomys*” becoming identified at the genus level as *Mus*).

To avoid issues with the variable reporting of pathogen classification varies by source, we harmonise pathogen names to ICTV binomial names. Where this is not possible we will retain the name used in the original study.

#### 4.4.2 Location of sampling

Locations were associated with administrative divisions using the Database of Global Administrative Areas (GADM) accessed through the `geodata R` package [59].

The coordinate resolution for coordinates labelled, site or study site will be set as 100 meters. For locations associated with an administrative area but no higher resolution data we record the coordinate resolution value as the radius of the administrative region to represent uncertainty in these coordinates.

#### 4.4.3 Imputed non-detections

Non-detection of a small mammal species in a location it may be expected to be (i.e., based on IUCN range maps or prior research) is of interest to researchers. We will enrich the detection-only data of included studies by imputing non-detections. This imputation will be restricted to species that have been detected at other sites or sessions in the study. Imputed non-detections will be labelled as imputed in the final data product.

### 4.5 Patient and Public Involvement statement

Individuals affected by Arenavirus or Hantavirus infections were not involved in the development of this study.

## 5 Discussion

This novel dataset will provide spatially and temporal information on small mammal sampling for Arenaviruses and Hantaviruses, offering a more comprehensive view of host-pathogen associations than is currently available. By including sampling effort and explicit spatial data, this dataset enhances the ability to quantify biases in where and how frequently sampling occurs, assess geographic gaps, and better understand the spatial ecology of these globally distributed pathogens of public health importance. The insights gained from this dataset could improve our understanding of how environmental changes, such as habitat fragmentation and urbanisation, influence pathogen dynamics and spillover risks.

Importantly, these data could inform the development of ecology-driven predictive models, helping to identify areas at heightened risk of zoonotic spillover [60,61]. The ability to link ecological data with human health outcomes could inform targeted public health interventions, enhancing outbreak preparedness and response. For instance, identifying hotspots of high pathogen prevalence in rodents could guide targeted development of ecologically-based rodent control measures in regions identified as vulnerable.

Existing host-pathogen datasets lack detailed spatial or temporal information, limiting their use in ecological and epidemiological modelling [62]. By addressing these gaps, our dataset provides information about host-pathogen interactions that span multiple spatial and temporal scales. Moreover, the explicit reporting of sampling effort allows for more robust analyses of detection probability, which is crucial for understanding the true distribution of pathogens [63,64].

While currently available global host-pathogen datasets do not account for sampling biases, the dataset proposed here will explicitly report the extent of sampling for pathogen within their host species, providing critical insights into spatial sampling biases and detection efforts. For instance, while it is expected that *Mastomys natalensis* sampling occurs throughout its range, the detection of specific pathogens such as LASV or Morogoro virus are currently only detected in their West African and East African radiation respectively. By detailing both presence and absence data, the dataset allows for a more nuanced understanding of pathogen distribution within host species, beyond mere occurrence records. Quantification of detection effort for specific pathogens is vital for assessing spatial sampling biases, identifying under-sampled regions, and refining ecological and epidemiological models [65].

Despite its strengths, this dataset has several limitations. First, the inherent variability in the quality and detail of reported data from included studies means that some records may lack critical information, such as exact coordinates or specific sampling dates. The imputation of non-detections, introduces assumptions that could affect the interpretation of absence data. Additionally, the dataset is inherently static will not reflect emerging data following production.

Furthermore, the reliance on published literature introduces publication bias, as studies with significant findings are more likely to be published than those with negative results [66]. This could skew the dataset towards areas and hosts with known or previously detected pathogens, potentially under representing regions where sampling has occurred but without positive detections. We will not routinely contact study authors to obtain data reported within publications. Therefore, data that are not reported within the article or its supplementary appendices are not included in the dataset. This is the subject of an ongoing data request study. To address limitations associated with extracting data without access to the raw data we encourage scientists with ongoing field studies and pathogen surveillance programmes, to submit their data to dynamic repositories (e.g., Pharos and GBIF [51,67]). Beyond research applications, the dataset could serve as a tool for policymakers, assisting in the identification of priority areas for viral discovery and public health interventions.

## 6 Conclusion

Overall, this dataset will be a valuable resource for understanding the ecology of Arenaviruses and Hantaviruses in their natural reservoirs. By bridging the gap between local-scale ecological studies and broader public health needs, it has the potential to enhance our ability to predict and mitigate the risks posed by these emerging pathogens. Continued efforts to update and expand this resource will be crucial for maintaining its utility in a rapidly changing epidemiological landscape.

## 7 Ethics and Dissemination

No ethical approval was sought. All data products of this project will be made available on GitHub (https://github.com/DidDrog11/arenavirus_hantavirus).

## 8 Authors’ contributions

D.S. - conceptualisation, methodology, data curation, investigation, software, supervision, writing - original draft, writing - review and editing R.R. - methodology, data curation, investigation, writing - review and editing H.G. - methodology, data curation, investigation, writing - review and editing A. M-C G. - methodology, data curation, investigation, writing - review and editing G. C. M. - methodology, supervision, writing - review and editing G. R. - methodology, data curation, investigation, writing - review and editing D. W. R. - funding acquisition, resources, supervision, writing - review and editing S. N. S. - conceptualisation, funding acquisition, methodology, resources, supervision, writing - review and editing

